# Relationships between migraine and interoception: normal accuracy but altered sensibility and behaviour

**DOI:** 10.1101/2024.11.11.622839

**Authors:** Siobhan Jones, Jessica Komes, Ester Benzaquén, Quoc C. Vuong, William Sedley

## Abstract

**Background:** Migraine is typically precipitated by unaccustomed changes in one’s internal state and/or external environment. Migraine symptoms largely involve increased, noxious, awareness of bodily changes and external stimuli. Links have been proposed between migraine and interoception (sensing and interpreting internal states), but direct evidence is limited.

**Methods:** Unmedicated, otherwise healthy, age-matched female participants were grouped by migraine tendency: control (no unprovoked headaches, n=19); low-frequency (<=3 migraines/month, n=20); high-frequency (>=4 migraines/month, n=19). Interoception was assessed, interictally, with standardised questionnaires such as MAIA-2 and a widely used heartbeat counting task.

**Results:** The notable significant questionnaire-based difference was in the ‘noticing’, ‘not distracting’ domain of the MAIA-2; controls were least likely to continue activities despite experiencing physical discomfort, and high-frequency migraineurs most likely. Follow-up questioning clarified that this behaviour related predominantly to migraine-related symptoms. The heartbeat task found no differences in accuracy, but lower confidence in the low-frequency migraine group than the control and high-frequency groups.

**Conclusions:** We suggest that low interoceptive confidence is a risk factor for migraine, whilst amplification of interoceptive signals caused by migraine restores this confidence, but at the price of migraine’s disabling symptoms. Self-reported tendency to deliberately ignore physical discomfort, including that caused by migraine, may result in more migraine attacks.

## Introduction

Migraine is a common yet incompletely understood nervous system condition, affecting approximately 12% of the population (1), with chronic migraine (CM) affecting 1-2%. It is likely that many more individuals with migraine are undiagnosed. In the UK around 10 million people aged 15-69 years old suffer with migraine (2), with an annual healthcare cost of £150 million.

Migraine is typically characterised by a headache, often severe, and a noxious hypersensitivity to physical or mental activity and external stimulation (for instance light or sound); it can be highly disabling and can feature nausea and vomiting (3). The migraine episode is divided up into four phases (not all of which are present in all individuals) comprising the premonitory phase characterised by autonomic and mood changes, (in some individuals) the aura phase characterised by focal neurological symptoms, the headache phase, and concluding with the post-drome phase (4).

Because migraine triggers largely consist of unaccustomed alterations in physiological state, we have previously posed the question of whether migraine helps to stabilise internal bodily states when these are pushed beyond predictable limits (5). Therefore, the question naturally arises of whether individual differences in people’s perception of internal states (interoception) are either of a cause and/or consequence of frequent migraines.

### Interoception

Interoception is the perception of sensations from inside the body (6), for instance heartbeat, respiration and hunger, and is vital for survival in informing the nervous system about the body’s physiological needs. It also encompasses the ability to interpret and respond appropriately to patterns of internal signals. Interoception interrelates with cognition, psychology and in particular, emotional regulation, and as such constitutes the afferent limb of allostasis (maintenance of physiological states across changing situations and requirements) (7). Interoceptive experience is influenced by both descending predictions about the expected state of the body and ascending visceral sensations (8)

### Measuring Interoception

Measurement of interoception can be divided into 3 domains: *interoceptive accuracy* (objective performance), *interoceptive sensibility (confidence)* (self-reported beliefs about the strength with which interoceptive signals are perceived) and *interoceptive awareness* (the correspondence between *accuracy* and *sensibility*; i.e. being more accurate when one thinks they are being more accurate, and vice versa) (9).

Sensibility is measured by questionnaires, or subjective self-report measures such as confidence performing an interoceptive task, whilst accuracy is based on objective task performance (10)

The ability to monitor one’s own heartbeats has emerged as a potential biomarker in some neurological and psychiatric disorders (22). Garfinkel et al, (2015) employed the heartbeat tracking task to determine individual differences in interoceptive accuracy. Research using heartbeat detection (HBD) has helped to shed light on key processes that relate to interoception.

### Links between Interoception and Migraine

#### Clinical overlap between migraine and interoception

Migraine is often considered within the wider context of physiological regulatory systems, i.e. homeostasis and allostasis, which rely on interoception as their main sensory input (5). Furthermore, between individuals, there is a strong correspondence between which sensory modalities act as triggers (e.g. hunger) and manifest premonitory symptoms (e.g. food cravings) (11), to the point that it can be difficult to assign a particular symptom to one category or the other (12).

#### Migraine and allostasis

Early concepts of homeostasis (maintaining rigid physiological stability) are inherently inefficient and have since been expanded and replaced by ‘allostasis’, defined as achieving “stability through change” (7). More specifically, allostasis describes the process whereby organisms change the defended levels of regulated parameters as needed to adjust to new or changing environments, including pre-emptive responses to anticipated changes (7).

Allostasis relies heavily on predictions, whose accuracy is indicated by the magnitude of ‘prediction errors’, which signal discrepancies between predicted and sensed physical states. For allostasis to function efficiently and safely, its underpinning predictions must be sufficiently accurate. We have recently characterised migraine as, in its ‘intended’ form, as an adaptive process to restore allostatic accuracy in the face of persistently elevated interoceptive prediction errors (IPE) (5). In this account, elevated IPE is the cause of migraine, amplification of IPE is the mechanism of migraine (leading first to the premonitory phase, then the migraine attack phase), and withdrawal from activity and stimulation is the forced behaviour that promotes stability to restore the accuracy of predictions. However, failure to respond to migraine episodes by reducing IPE, or external or internal environments that present ongoing unavoidable sources of error, is thought to lead to persistent, recurrent, and self-reinforcing amplification of IPE, clinically causing high-frequency episodic or chronic migraine, and physiologically leading to excessive stress responses that result in a kind of permanent damage termed ‘allostatic load’ (13).

#### Migraine as a condition of interoception

The idea that migraine is primarily a condition of altered interoceptive processing, which sits well within its existing categorisation as “a disorder of sensory processing” (3). Migraine may also be a phenomenon that straddles multiple categories, being adaptive in some instances and disordered in others (much like pain) (8, 24). Interoceptive abilities could therefore be a key factor in determining the susceptibility to, and the frequency/chronicity of, migraine. For instance, greater interoceptive accuracy might help to prevent migraines by reducing IPE themselves, and/or through the formation of more accurate predictions of future physiological states (5). Additionally, or alternatively, greater interoceptive sensibility (the subjective strength of interoceptive signals) could result in fewer migraines (if IPE are highlighted and addressed earlier) or more migraines (if amplified IPE cross the threshold to trigger migraines more often). Furthermore, behavioural responses to interoceptive signals may be important, for instance whether an individual acts on or ignores a particular sensation, and this in turn may depend upon metacognitive interoceptive awareness. Finally, in the opposite direction of causation, migraine episodes might cause lasting alterations in interoception, for instance as a partial carryover of the noxiously increased interoceptive signals experienced during migraine attacks.

### Aims and hypotheses

Migraine and interoception is a largely unexplored area but its importance is starting to be recognised(14). We conducted a study to ascertain abnormalities of interoception as a predictor of migraine frequency by employing a Heartbeat Tracking Task and objective and subjective measures of interoception in the form of questionnaires in order to capture the three interoceptive domains.

## Methods

### Participants

We recruited age-matched groups of female participants aged between 18 and 65, stratified by self-reported migraine frequency, into Low Frequency (LF), High Frequency (HF) and Control groups (Ctrl). The LF group comprised 20 participants with LF migraine (≤=3 migraine days per month), the HF group comprised 19 participants with HF migraine (≥= 4 migraine days per month), and the Ctrl group comprised 19 participants with no history of unprovoked headaches (even reports of ‘tension headache’ was an exclusion criterion in case it reflected a limited or milder form of migraine). The threshold of 4 migraine days per month was chosen to give the most bimodal distribution of monthly migraine days across screened participants. It also accords with the typical threshold at which prophylactic treatment would be offered. Ethical approval was provided by Newcastle University (Ref: 37672/2024). Informed written consent was obtained from all participants. All methods were performed in accordance with institutional procedures and guidelines, and with General Data Protection Regulation (GDPR), Good Clinical Practice (GCP) for research and the Declaration of Helsinki.

### Case history and questionnaires

Once screened for eligibility and enrolled, participants were sent the following validated mood and behavioural questionnaires via an online link to complete prior to attendance for laboratory testing:

- Body Perception Questionnaire (BPQ) short form (16)
- Migraine Specific Quality of Life Questionnaire (MSQL) (17)
- Hospital Anxiety and Depression Scale (HADS) (18)
- Intolerance of Uncertainty Scale (IUS) (19)
- Multidimensional Assessment of Interoceptive Awareness (MAIA-2) (20)
- Inventory of Differential Interoceptive Awareness-Short form (IDIA) (21)

On attendance in person, the lead researcher took a detailed case history to confirm the diagnosis of migraine according to the ICHD-3 criteria (22) and estimate the number of monthly migraine days, headache days, and presence or absence of aura.

The experiment was conducted on interictal days (defined as more than 24 hours since most recent headache, to avoid postdrome changes, and more than 24 hours prior to subsequent migraine, to avoid inadvertently capturing premonitory changes). Because of this requirement, an exclusion criterion was headache or migraine symptoms ‘on most days’, to the point where satisfying the interictal criteria would be difficult. As these might in principle affect interoception, participants were excluded who were taking any migraine preventative medication, or any regular medication with a known action on the central, peripheral, or autonomic nervous system, including sedatives, antidepressants, antihypertensives and anticonvulsants. Other exclusion criteria included any known abnormality of brain structure (e.g., stroke, tumour), or other neurological disorder (e.g., multiple sclerosis or epilepsy), psychiatric comorbidity of sufficient severity to limit certain activities of everyday life, or abnormality of cardiac rhythm or function. Participants were also advised not to attend if they had taken a triptan in the last 24 hours. The Ctrl group criteria included not experiencing headaches ever except during physical illness. The rest of the exclusion criteria were the same as the migraine participants. In the migraine groups the participants were also asked to notify the researcher if they suffered a migraine 24 hours after the testing. The protocol was to exclude data from these participants, but no participants reported a migraine in the 24 hours after testing.

### Heartbeat tracking task

Participants performed the heartbeat tracking task, implemented in Matlab (23) using the Psychtoolbox toolbox (24), in a dimly lit sound-attenuated room (10). Participants were asked to count heartbeats by only concentrating on their body and not by taking their pulse or using any exteroceptive cues. The task simply required them to count the number of heart beats occurring during a specified time interval. There were 20 trials (with 1 extra practice trial), which differed in length, in randomised order, and were indicated with auditory stimuli (“Start” and “Stop”) presented with free-field speakers. Four repetitions of each of the following durations were included: 17s, 22s, 27s, 37s, 47s. At the end of each trial participants were asked the on-screen question ‘How many beats did you count?’, followed by a second question of ‘How confident are you?’ (in their response being accurate) and given a scoring scale between 0 and 100 with ‘0’ equating to not confident, and with 100 being very confident (10). The reported number of beats was then compared to the actual number of beats as extracted from the ECG monitor using a Biosemi ActiveTwo system (25). ECG was recorded from the left wrist, and heartbeats counted using the Pan-Tompkin algorithm (26), with manual confirmation of accuracy of fit for each trial. Accuracy and awareness scores were measured by examining the relation between the participants perception (self-reported account of their heartbeats) in comparison with the actual recorded heartbeats. Analysis was performed off-line, and participants were not given feedback about their performance. The accuracy was calculated as 1 - |a-p| / ((a+p)/2) where ‘a’ is the actual number of heartbeats, and ‘p’ is the perceived number of heartbeats, i.e. one minus the absolute difference between actual and perceived number of beats divided by the mean of the actual and perceived number of beats. Awareness was calculated by the strength of linear correlation, across trials, between accuracy and confidence, expressed as Pearson’s r.

### Statistical Analysis

All analyses were performed using the SPSS software (27). One-way ANOVA was used to detect differences between participant groups, followed by pairwise T tests as post-hoc tests in cases where there was a main effect of group.

Due to the high number of total questionnaires and their domains, and substantial overlap between these, we used a Principal Component Analysis (PCA) to identify the common underlying factors to the questionnaires. An eigenvalue threshold of 1 was used to select a final number of components for analysis. Components were related to questionnaire scores and scales via a rotated matrix of loadings, and a threshold of +/− 0.4 was used to indicate a meaningful association between each component and score/subscale. Statistical analysis was run on principal components for questionnaire data, and on individual measures (accuracy, confidence and awareness) for the heartbeat tracking task.

## Results

Subject characteristics are summarised in Table 1.

**Table 1:**
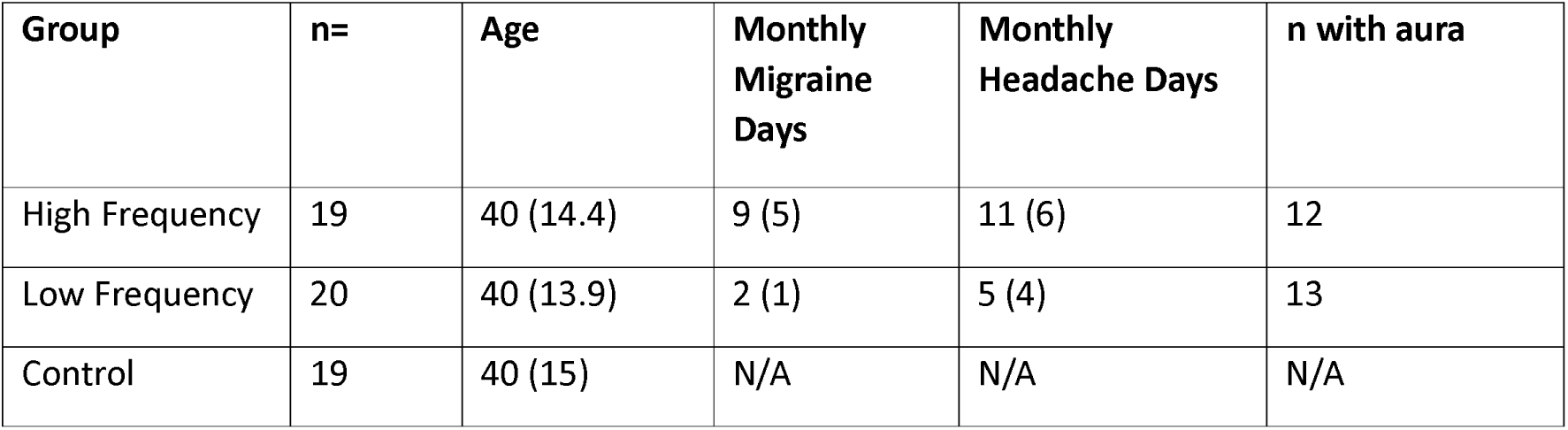
Group characteristics. Fifty-nine age-matched female participants were grouped by migraine tendency: control (no unprovoked headaches, n=19); low-frequency (<=3 migraines/month, n=20); high-frequency (>=4 migraines/month, n=19). Convention for numbers = mean (SD).

### Heartbeat tracking task results

The one-way ANOVA analyses for main effect of group (Figure 1A) showed no difference in interoceptive accuracy (F = .293, p = .747), whilst interoceptive confidence (Figure 1B) showed a significant difference (F = 3.512, p = .037), and interoceptive awareness (figure 1C) showed a non-significant trend (F =.294, p =.746) which visually resembled the relative group differences in confidence. For confidence, the Bonferroni-corrected post-hoc comparisons did not quite reach statistical significance (Ctrl vs. HF p = 1.000, HF vs LF p = .103, LF vs Ctrl p = .062), though the clear impression was that the LF group scored lower than the Ctrl and HF groups.

**Figure 1:**
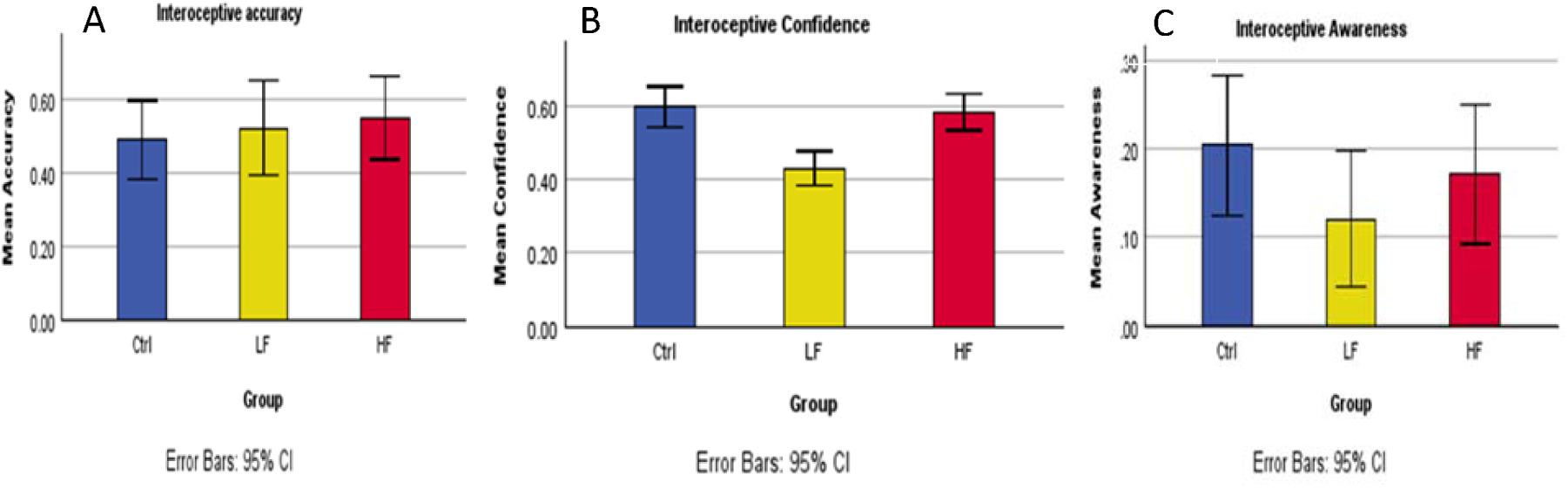
Results of the heartbeat counting task. Interoceptive accuracy (A) showed no significant effect of group, with only a weak trend towards higher accuracy in migraine groups. Interoceptive confidence (B) showed lower confidence in the LF group than the Ctrl groups. Interoceptive awareness (C) showed a non-significant trend.

### Questionnaire responses

#### Component structure

The component analysis revealed 5 components, whose eigenvalues are displayed in Table 2, and matrix of (rotated) loadings in Table 3. Based on their loadings, we interpret the component structure as follows, and is demonstrated in Figure 2: Component 1 was a general interoceptive abilities measure, showing no effect of group (F =.285, p =.753); Component 2 related to intolerance of uncertainty, anxiety, worry, and physical sensations associated with these, and showed a near-significant group effect (F = 2.999, p = .058). Component 3 indicated cardiorespiratory sensations, and showed no effect of group (F = .096, p = .908); Component 4 indicated depression, and showed no effect of group (F = .462, p = .632); Component 5 related to noticing more bodily discomfort, and modifying activities less in response to discomfort, and demonstrated a significant main effect of group (F = 3.943, p = .025). A Bonferroni-corrected post-hoc analysis on Component 5 showed that the HF group scored higher than Ctrl group (p = 0.023), whilst there was no significant difference between the LF and Ctrl groups (p = .995), and only a trend between the HF and LF groups (p = .208).

**Table 2:** Rotation Sums of Squared Loadings.

| Component | Eigenvalue | % Variance | Cumulative<br>% Variance |
| --- | --- | --- | --- |
| 1 | 3.602 | 24.014 | 24.014 |
| 2 | 2.384 | 15.896 | 39.911 |
| 3 | 2.041 | 13.610 | 53.521 |
| 4 | 1.568 | 10.451 | 63.971 |
| 5 | 1.367 | 9.111 | 73.082 |

**Table 3:** Component Matrix. Table of rotated loadings of components onto questionnaire domains. Values exceeding +/− 0.40 are displayed, indicating ‘meaningful’ associations. Negative values indicate a higher value of that component for an individual indicating a lower score on that questionnaire (domain).

| Component | 1 | 2 | 3 | 4 | 5 |
| --- | --- | --- | --- | --- | --- |
| BPQ: Body Awareness |  |  | .648 | -.499 |  |
| BPQ: Supradiaphragmatic Reactivity |  |  | .803 |  |  |
| IUS |  | .651 | .866 | -.497 |  |
| MAIA2: Noticing | .624 |  |  |  | .524 |
| MAIA2: Not distracting |  |  |  |  | -.851 |
| MAIA2: Not worrying |  |  |  |  |  |
| MAIA2: Attention regulation | .774 | -.730 |  |  |  |
| MAIA2: Emotional Awareness | .773 |  |  |  |  |
| MAIA2: Self-regulation | .859 |  |  |  |  |
| MAIA2: Body Listening | .722 |  |  |  |  |
| MAIA2: Trusting | .739 |  |  |  |  |
| IDIA |  | .701 |  |  |  |
| HADS-A |  | .790 |  |  |  |
| HADS-D |  |  |  | .809 |  |
| Component interpretation | General interoceptive abilities | Anxiety | Cardiorespiratory / gastrointestinal sensations | Depression | Noticing and not responding to discomfort |

**Figure 2:**
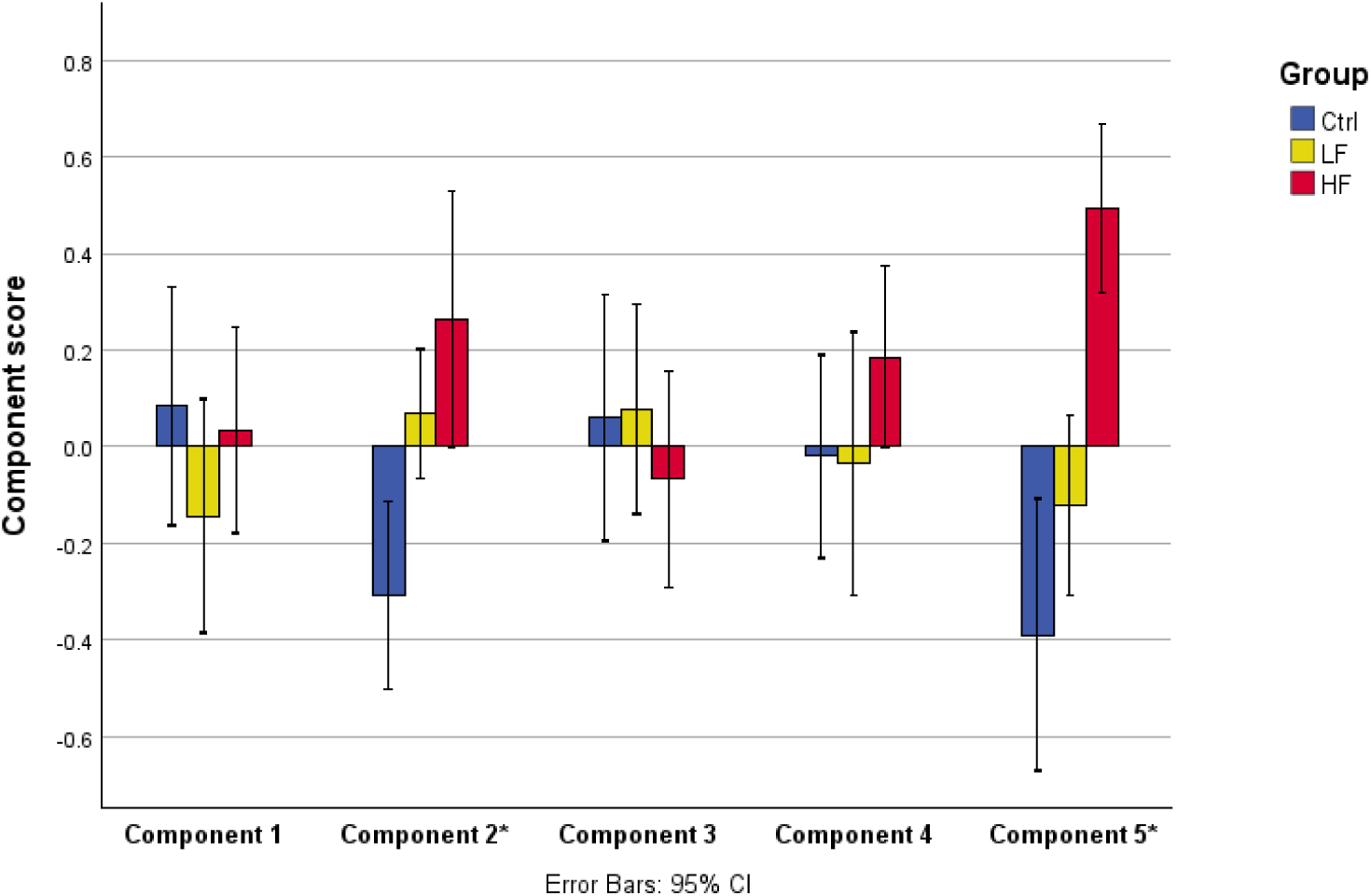
Group comparison of scores on the five questionnaire components. Asterisks indicate components either showing a significant effect of group (Component 5), or a near-significant effect deemed probably meaningful on account of strong accordance with prior literature (Component 2). Note that higher scores for Component 5 indicate noticing more discomfort and reducing or modifying activity less on account of discomfort.

#### MAIA-2 Subscales (Component 5)

To determine whether differences in Component 5 scores related to migraine-related or non-migraine-related symptoms, we re-tested identical questions of the ‘noticing’ domain e.g. (“when I am tense I notice where the tension is located in my body”) and ‘not-distracting’ domain e.g. (“I distract myself from sensations of discomfort”) of the MAIA-2 to the migraine groups in two ways, using identical questions, but with two separate frames of reference, indicated by a prefacing instruction:

1. “Relating to migraine symptoms only…”
2. “Relating to non-migraine symptoms…”

The Ctrl group were re-tested the questions without this differentiation, to maintain parity.

These scores were multiplied by the component loadings indicated in Table 3, and Eigenvalue in Table 2, to yield ‘projected Component 5’ scores (which were similar but not identical to original Component 5 scores, on account of not incorporating loadings less than +/−0.40). These are displayed in Figure 3. The follow-up questionnaires were only answered by a minority of participants (9 HF, 14 LF and 15 Ctrls).

**Figure 3:**
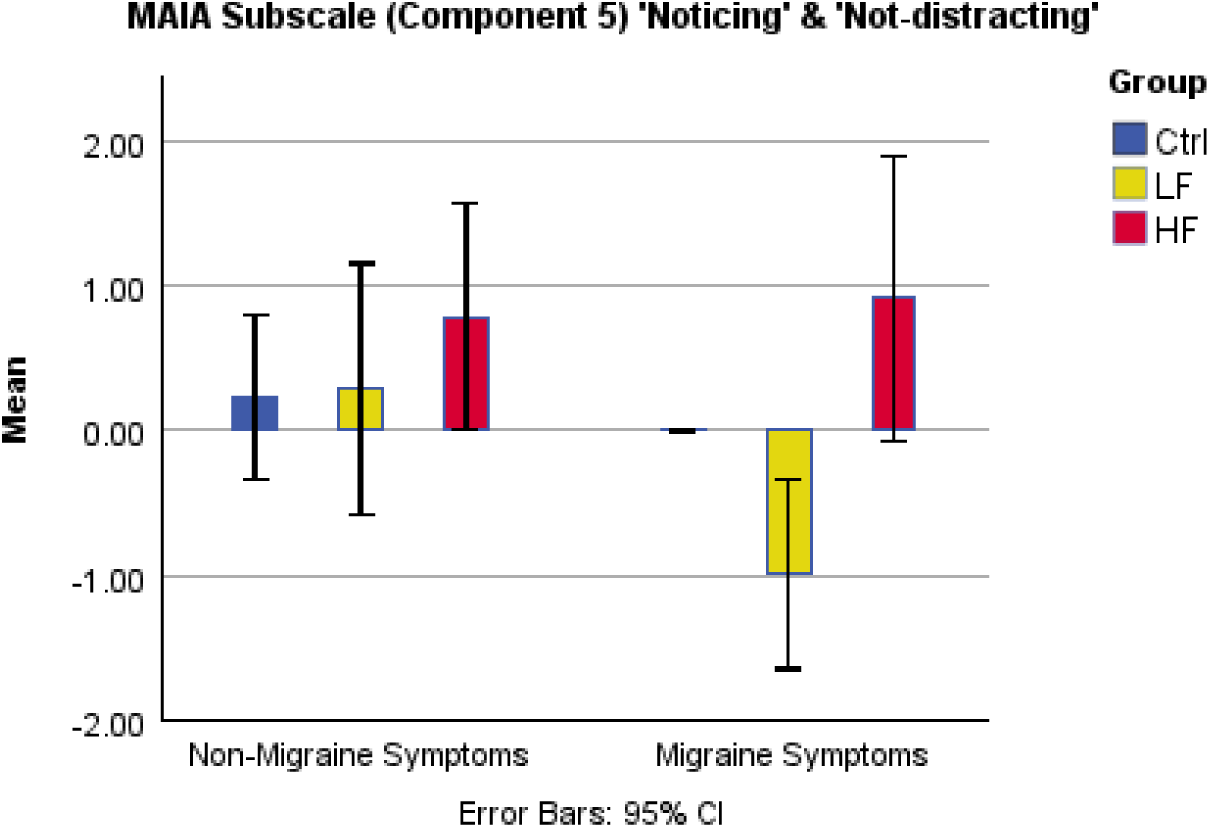
Projected Component 5 (MAIA Noticing and Not Distracting domains) scores specifically relating to non-migraine and migraine symptoms. Controls, by definition, have a score of zero for migraine symptoms.

One way ANOVA referring to ‘non-migraine’ symptoms demonstrated non-significant main effect of group F= .711, p = .498.

Independent t-tests comparing Projected Component 5 scores between HF group and LF groups referring to migraine symptoms showed higher scores in the HF group (t(21) = 4.150, p = <.001), and between HF group and LF group referring to non-migraine symptoms showed t(21) = .925, p = .365.

## Discussion

### Interoceptive confidence is reduced in low-frequency migraine and restored in high-frequency migraine

Interoceptive accuracy scores were not reduced in migraine participants. This displays a stark contrast with reported findings of interoceptive accuracy in people with chronic pain that is both lower than controls, and negatively correlate with symptom severity (27). In the present study, interoceptive confidence was lower in LF, and normal in HF. We speculate that in the LF migraine group, low interoceptive confidence (sensibility) (10) is a risk factor for migraine, whilst lasting amplification of interoceptive signals caused by migraine restores this confidence, but at the price of migraine’s disabling symptoms. In the HF group this would then indicate the ‘effect’ of migraine, i.e. acting to increase interoceptive sensibility.

### Tendency to notice and ignore physical discomfort is associated with higher migraine frequency

Questionnaire responses indicated that the HF migraine group reported a tendency to notice discomfort more than the other groups (‘noticing’ subscale of MAIA2) but modify or reduce their activity less as a result (‘not distracting’ subscale of MAIA2).

Clarification revealed that this was predominantly driven by migraine symptoms, though a similar but weaker trend was observed for non-migraine-related symptoms. A proportion of these scores likely reflected living with an increased burden of migraine symptoms, particularly the ‘noticing’ domain, whereas the larger ‘not-distracting’ contribution to the component is harder to fully explain in such a way. It is equally possible that the tendency to ignore or ‘push-through’ migraine-related symptoms hinders the utility of migraine to resolve interoceptive errors. If we accept the notion that the ‘mechanism’ of migraine is to increase the strength of interoceptive signals (i.e. interoceptive sensibility) in order to enforce behaviour that resolves errors, but these signals are then ignored by the individual continuing activities despite feeling bodily discomfort, then this might lead to even more migraines as a protective reflex (and a positive feedback loop ensuing) as illustrated in Figure 4. This notion aligns well with the existing concept of ‘error awareness’ in migraine, in which repeated migraine episodes lead to a lasting increase in the awareness of interoceptive errors, which become aversive on account of being associated with migraine symptoms (28).

**Figure 4:**
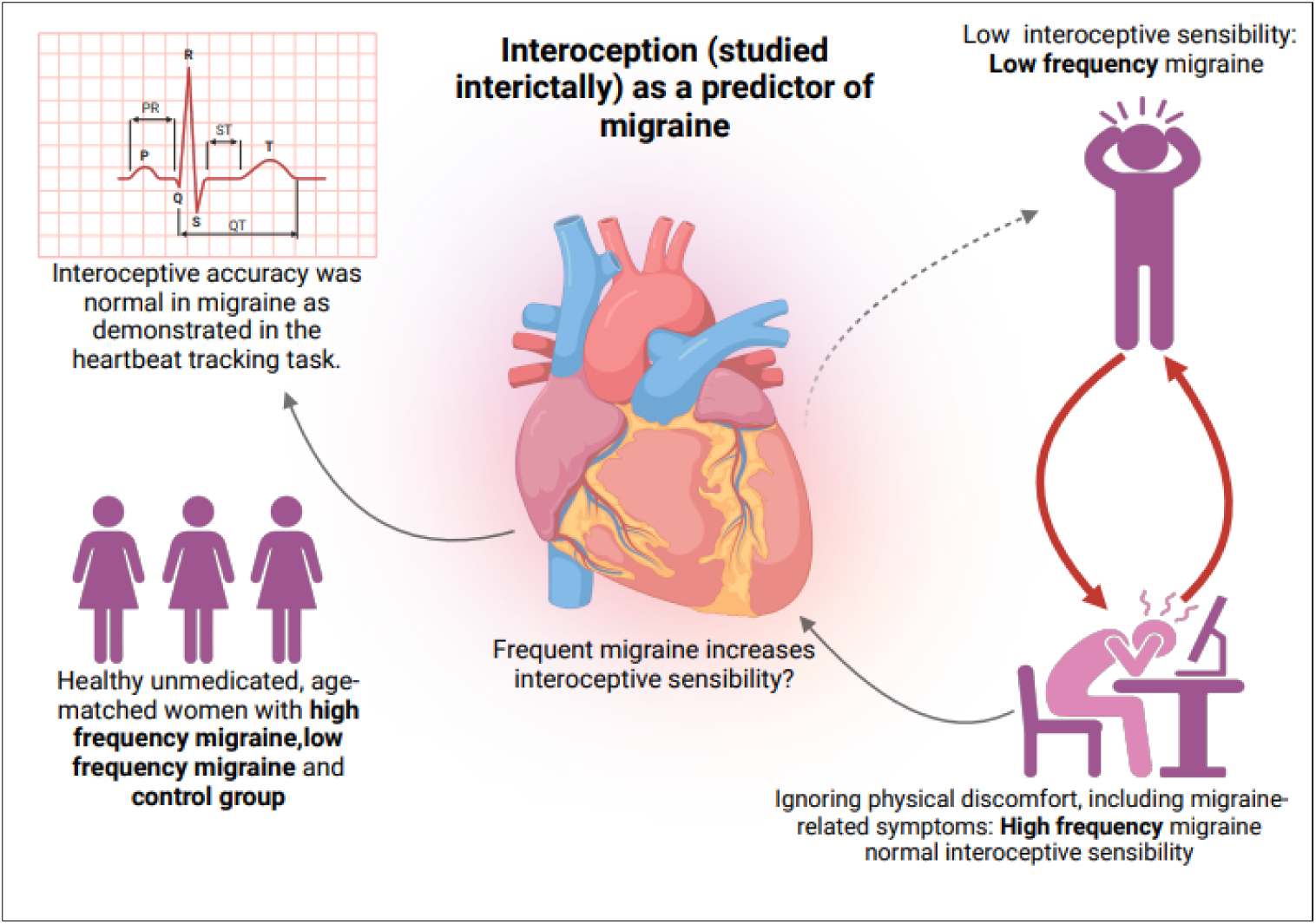
Schematic summary of interoception (studied interictally) as a predictor of migraine.

### Strengths, limitations and future directions

This has been a specific study on ‘ordinary’ migraine, unconfounded by sex/gender, medication, or physical or mental health comorbidity. It has allowed us to be able to isolate only interictal differences, to avoid confounding immediate effects of migraine processes. We excluded chronic migraine and the more complex patients typical of tertiary headache clinics, where migraine is more ‘disordered’.

Limitations of the study included the relatively low number of participants, examining only one interoceptive domain objectively, and not longitudinally examining the immediate and short-term effects of migraine on interoception. One must take into consideration the measures of interoceptive ability and how they may differ due to required cognitive effort, and on the nature, length and the difficulty of the task (29). Assessing separate interoceptive domains such as respiration could help address technical concerns, and search for generalisation of findings (30)

Future directions could look at chronic migraine and also how migraine interrelates to commonly co-morbid conditions, such as chronic pain, in which interoceptive accuracy deficits have been observed (31) This could also include longitudinal tracking over a migraine cycle, to assess putative claims more directly about ‘causes’ and ‘effects’ of migraine.

Our results also highlight the potential insights possible through questionnaire/subjective measures of interoception, not all relying on lab-based tests of accuracy, which opens the door to large-scale online questionnaire studies and incorporation into clinical consultations. Longer-term aims are to use this information to develop innovative interventions such as interoceptive training programmes to help prevent the conditions that cause migraine episodes from occurring.

## Conclusions

These data offer insight into the role that migraine tries to play in helping us safely regulate our physiology, lending some empirical support to our recently proposed model of migraine as a ‘failsafe’ to stabilise homeostasis/allostasis, and may help lead to a conceptual framework for patients to help them understand and non-pharmacologically manage their migraine by addressing its underlying ‘causes’. This study has provided some new interesting and theoretical findings in the relationship between interoception and migraine and has highlighted some potentially fruitful avenues for further research.

***The author(s) disclosed receipt of the following financial support for the research, authorship, and publication of this article: This work was supported by the The Wellcome Trust [222071/Z/20/Z]* .**

## Data Availability

The datasets used and/or analysed during the current study is available from the corresponding author on reasonable request.

